# Pan-viral ORFs discovery using Massively Parallel Ribosome Profiling

**DOI:** 10.1101/2023.09.26.559641

**Authors:** Shira Weingarten-Gabbay, Matthew R. Bauer, Alexandra C. Stanton, Susan Klaeger, Eva K. Verzani, Daniel López, Karl R. Clauser, Steven A. Carr, Jennifer G. Abelin, Charles M. Rice, Pardis C. Sabeti

## Abstract

Unveiling the complete proteome of viruses is crucial to our understanding of the viral life cycle and interaction with the host. We developed Massively Parallel Ribosome Profiling (MPRP) to experimentally determine open reading frames (ORFs) in 20,170 designed oligonucleotides across 679 human-associated viral genomes. We identified 5,381 ORFs, including 4,208 non-canonical ORFs, and show successful detection of both annotated coding sequences (CDSs) and reported non-canonical ORFs. By examining immunopeptidome datasets of infected cells, we found class I human leukocyte antigen (HLA-I) peptides originating from non-canonical ORFs identified through MPRP. By inspecting ribosome occupancies on the 5’UTR and CDS regions of annotated viral genes, we identified hundreds of upstream ORFs (uORFs) that negatively regulate the synthesis of canonical viral proteins. The unprecedented source of viral ORFs across a wide range of viral families, including highly pathogenic viruses, expands the repertoire of vaccine targets and exposes new cis-regulatory sequences in viral genomes.

## INTRODUCTION

Despite remarkable advances in sequencing viral genomes, functional annotations of these genomes have lagged behind, hindering our understanding of the mechanisms by which viruses propagate and interact with the immune system. One important area of exploration is the translatome – the entire collection of viral mRNA sequences being translated into proteins. Beyond the annotated canonical ORFs, viral genomes encode non-canonical ORFs that do not fulfill the classical definition of ORFs, i.e. do not start with an ATG codon and/or are shorter than 100 amino acids (aa). These non-canonical ORFs and the resulting microproteins have been shown to play a critical role in modulating viral infection (Lulla et al. 2019; Ogden et al. 2019; Lulla and Firth 2020), contributing to the immune response to viruses (Bansal et al. 2010; Ingolia et al. 2014; Yang et al. 2016; Weingarten-Gabbay et al. 2021), and regulating the expression of canonical viral proteins (A. Chen, Kao, and Brown 2005; Gould and Easton 2005; Shabman et al. 2013). However, the deviation from classical ORF features challenges their detection computationally and relies mostly on experimental measurements. Thus, for decades of research this “hidden” source of proteins has remained mostly unknown.

The development of ribosome profiling over the past decade has transformed our ability to detect translated regions across genomes (Ingolia et al. 2009). Ribosome profiling (Ribo-seq) utilizes deep sequencing of ribosome-bound mRNA fragments to determine ribosome occupancy, indicating translated regions at single nucleotide resolution. Quickly adapted to different organisms, ribosome profiling uncovered a striking number of non-canonical ORFs in mammalian cells, yeast, bacteria and viruses including upstream and upstream-overlapping ORFs (uORFs and uoORFs) in 5’UTRs, short ORFs in non-coding RNAs, and overlapping internal ORFs in annotated coding sequence (iORFs) (Ingolia, Lareau, and Weissman 2011; Ingolia, Hussmann, and Weissman 2019).

Although undoubtedly important, the landscape of translated regions is still unknown for the majority of viruses. Ribosome profiling has been successfully employed to profile a handful of viruses, however, it is still limited in throughput with respect to the number of viruses that can be tested. Each virus requires a unique culturing system, exhibiting selective tropism to cells and growing conditions. Thus, each virus typically needs to be profiled in isolation, limiting the number of viruses that can be assayed in each experiment. In addition, researching highly pathogenic viruses necessitates high-containment facilities and strict safety protocols that challenge the execution of ribosome profiling within biosafety level 3 or 4 (BSL3/4) laboratories. Moreover, some viruses cannot be cultured in the laboratory, making it impossible to profile their translation *in vitro*. Finally, since viruses are a moving target, their genome changes constantly and we can benefit from a method that will evaluate many variants in parallel.

Creating an MPRP platform for the discovery of ORFs across many viral genomes, independent of their culturing conditions and biosafety level, would transform our knowledge of viruses’ translatomes and is poised to expose new biology. In this study, we leveraged the power of a fully designed oligo synthesis library (Weingarten-Gabbay et al. 2016, 2019; Seo et al. 2023) and combined it with ribosome profiling to perform pan-viral ORF discovery. We measured the translation of 20,170 synthetic sequences from 679 viral genomes in two human cell lines, resulting in the identification of thousands of new ORFs. We estimated ORF discovery using the annotated CDSs in these genomes and non-canonical ORFs that were identified in traditional ribosome profiling of infected cells. We then examined the function of the new ORFs that we detected in two processes: HLA-I antigen presentation and uORF-mediated translation regulation of canonical viral proteins.

## RESULTS

### Developing Massively Parallel Ribosome Profiling to identify ORFs in 679 viral genomes

To measure ribosome occupancy and infer translated regions across hundreds of human viruses, we developed Massively Parallel Ribosome Profiling (MPRP) (**Figure 1A**). Instead of infecting cells with individual viruses, we used oligonucleotides library synthesis technology to encapsulate thousands of viral sequences in a single pooled experiment. Each oligo contained 200nt of viral sequence, flanked by constant primers. We amplified the library using the constant primers and cloned it into an overexpression plasmid downstream of a CMV promoter and upstream of a Woodchuck Hepatitis Virus (WHV) Posttranscriptional Regulatory Element (WPRE) used to further enhance expression. To monitor the translation of ORFs in the designed oligos, we excluded ATG start codons on the plasmid and the constant primer sequence upstream to the cloned oligo. Since the designed sequence is limited to 200nt, some oligos contain the beginning of a viral ORF but not the end. We ensured translation termination in the absence of the endogenous stop codon by inserting stop codons in all three potential reading frames on the plasmid, downstream of the cloned oligo. We transfected the pooled library plasmid into HEK293T human embryonic kidney cells or A549 human lung cells. We then performed a modified protocol of ribosome profiling on library-transfected cells (McGlincy and Ingolia 2017) after treating cells with either cycloheximide (CHX) to inhibit elongating ribosomes across the entire ORF, or lactimidomycin (LTM) to specifically inhibit initiating ribosomes at the start codon. We then mapped deep sequencing reads representing ribosome footprints to the synthetic library and identified ORFs using PRICE, a computational method for the detection of canonical and non-canonical ORFs in ribosome profiling experiments (Erhard et al. 2018).

**Figure 1.**
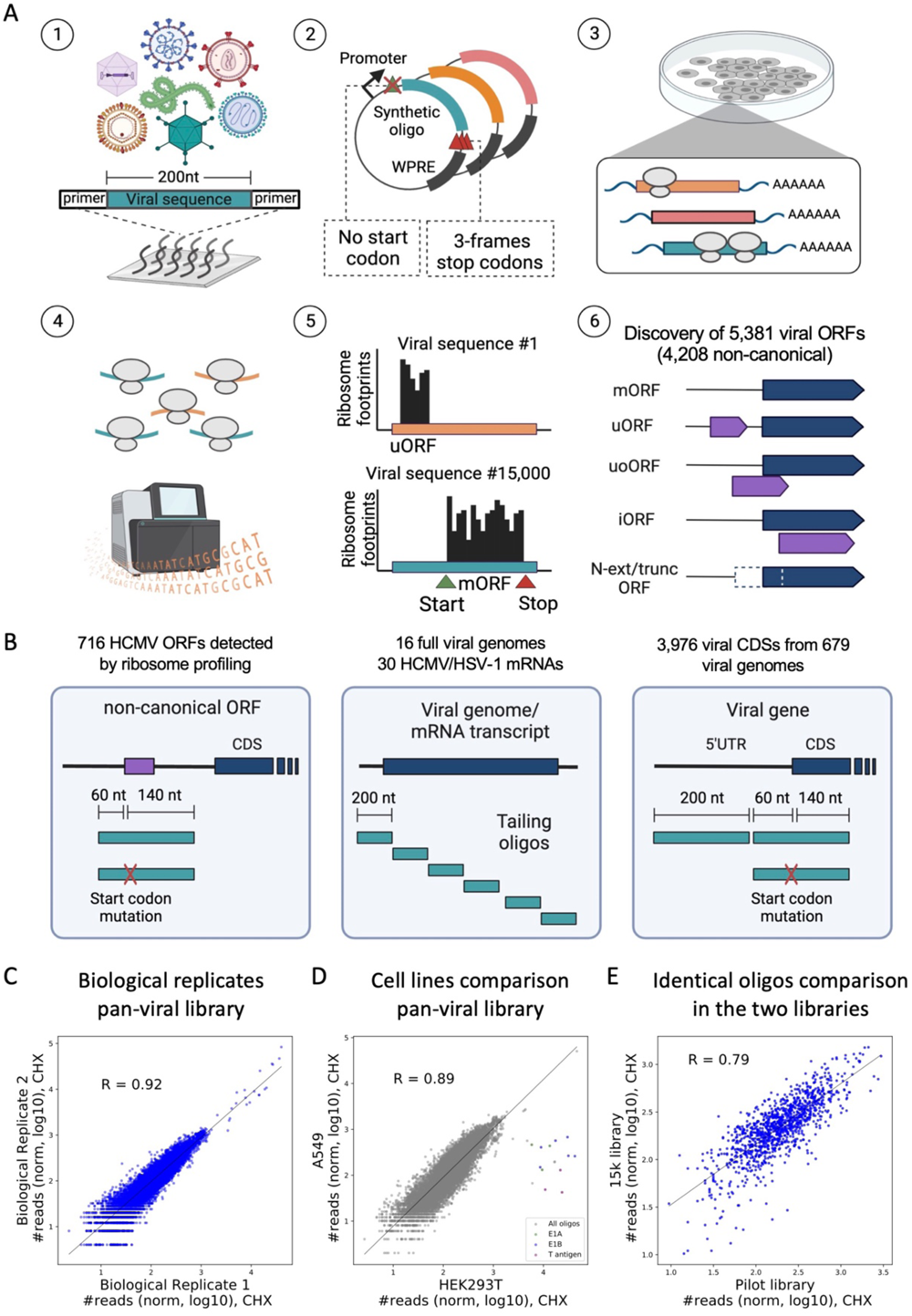
Design of oligonucleotide synthetic library and MPRP measurements. **(A)** Illustration of the Massively Parallel Ribosome Profiling experiment (MPRP). (1) Synthetic library amplification using constant primers. (2) Cloning library into overexpression vector. (3) Transient transfection of plasmid pool into HEK293T or A549 cells for 24 h. (4) Treating cells with either LTM or CHX and performing ribosome profiling protocol. (5) Mapping deep sequencing reads (representing ribosome footprints) to the synthetic library. (6) Inferring translated ORFs using PRICE (Erhard et al. 2018). **(B)** Design of the tested synthetic oligonucleotides: (i) ORFs that were identified by ribosome profiling in infected cells with either intact start codon or a GCC mutation. (ii) Tailing oligos encompassing complete viral genomes/transcripts. (iii) Oligos spanning the 5’UTR and the first 140 nt of annotated viral CDSs. For the region containing the CDS, two oligos were designed: the wild type sequence and a start codon mutated oligo. **(C)** Comparing the number of ribosome footprints mapped to 15,000 oligos in two biological replicates of MPRP experiment in HEK293T cells. R=0.92, Pearson correlation. **(D)** Comparing the number of ribosome footprints mapped to 15,000 oligos in the MPRP experiment in HEK293T and A549 cells. R=0.89, Pearson correlation. **(E)** Comparing the number of ribosome footprints mapped to 1,163 identical oligos in two synthetic libraries that were independently cloned, transfected and measured using MPRP in HEK293T cells. R=0.79, Pearson correlation.

We initially verified that our experimental system captures the translation of an ORF embedded within a synthetic oligo. We performed ribosome profiling on cells that were transiently transfected with a full-length Green Fluorescent Protein (GFP) (**Figure S1A**) and observed the expected ribosome footprints on the GFP CDS (**Figure S1B**). We then swapped the full-length GFP with the first 170nt of the reporter that resembled the length of a synthetic oligo in the library. Although the cloned sequence contained only part of the original ORF and did not harbor the GFP stop codon, we detected ribosome footprints on the truncated GFP regions, with successful termination in one of the three stop codons that we added to the plasmid (**Figure S1C**). Interestingly, we observed unexpected ribosome footprints on the WPRE region. These footprints accounted for a large fraction of the sequencing reads in the MPRP experiment because they originated from a constant region of the overexpressed plasmid. To remove WPRE-derived footprints prior to sample sequencing, we implemented a homemade ribosomal RNA (rRNA) depletion step (Methods) and appended WPRE-targeting probes in addition to the rRNA probes.

We designed a first ‘pilot’ library of 5,170 oligos to estimate viral ORF discovery using MPRP. This library contained 716 ORFs that were identified in HCMV-infected cells by traditional ribosome profiling (Stern-Ginossar et al. 2012) (**Figure 1B**). Each oligo was composed of 60 nt upstream of the start codon and 140 nt downstream of the start codon. In addition, for each ORF, we also designed a mutated oligo in which we replaced the reported start codon with GCC. To evaluate de novo discovery of ORFs, we tiled the entire genome of 16 viruses, and 30 mRNAs annotated in the genomic datasets of HCMV and HSV-1 (NC_006273.2 and JN555585.1, respectively) (**Figure S2A**). When examining the number of ribosome footprints across oligos from different regions of the mRNA transcripts, we observed the expected low occupancy in the 5’UTR and enrichment of ribosomes at the start codon (**Figure S2B**). However, we also observed high occupancy along the CDS. Since most of the 200 nt-long oligos tailing the CDS region do not contain the main CDS start codon, ribosomes in these oligos represent independent translation initiation from an alternative start codon downstream of the CDS start codon. However, in the context of the full transcript, these start codons are preceded by multiple start codons, making it unlikely that the ribosome will initiate at this position. We reasoned that MPRP has higher accuracy at the beginning of annotated CDSs and 5’UTRs, which better mimic the native genomic context. Thus, we decided to focus on these regions for the design of the pan-viral library.

We designed a second ‘pan-viral’ library of 15,000 oligos to screen for novel ORFs in the 5’UTRs and the beginning of the CDS of 3,976 genes in 679 viral genomes. For each gene we designed three oligos: a wild type oligo containing 60 nt of the 5’UTR and 140nt of the CDS, a mutated oligo in which the annotated start codon was mutated to GCC, and a farther upstream oligo in the 5’UTR region, starting -260 nt relative to the CDS start codon (**Figure 1B**).

To gauge the reproducibility of the MPRP measurements, we compared ribosome occupancy on the synthetic oligos in different experiments. Measurements of the pilot and the pan-viral libraries were consistent between biological replicates in HEK293T cells (R=0.81 and R=0.92, respectively, **Figure S3 and** **Figure 1C**). We also found high agreement between ribosome occupancies on the pan-viral library in HEK293T and A549 cells (R=0.89, **Figure 1D**). A small number of oligos had higher occupancy in HEK293T than A549 cells; closer examination of these outliers reveal shared sequences with the adenoviral E1A/B genes and the SV40 large T-antigen that are endogenously expressed by HEK293T cells (Tan et al. 2021). Thus, the elevated occupancies likely stem from cross-mapping of endogenous ribosome footprints to the synthetic library. Finally, we designed a set of 1,033 oligos with identical sequences in both the pilot and pan-viral libraries and found good agreement between oligos that were independently synthesized, cloned and measured using MPRP (R=0.79, **Figure 1E**). Altogether, these data indicate that MPRP produces reproducible ribosome occupancy measurements on thousands of viral sequences.

Our MPRP measurements on the pan-viral library uncovered 5,381 viral ORFs including 4,208 non-canonical ORFs (uORFs, uoORFs, iORFs, N-extended isoforms, and N-truncated isoforms) (Table S1) for further study.

### Estimating ORF discovery using annotated viral coding sequences

To estimate the detection of viral ORFs when expressed from a synthetic library, we examined the distribution of ribosome footprints across oligos representing the annotated CDSs (Figure 1B). We performed metagene analysis for all the CDSs from each family by computing the average ribosome footprints in each position relative to the annotated start codon (Figure 2A). As expected, we found enrichment of ribosome footprints in the CDS region with higher density at the beginning of the sequence, a phenomenon often observed in ribosome profiling measurements (Figures 2A**, 2B**). Since ribosome profiling provides single nucleotide resolution of ribosome footprints, it can be used to infer the reading frame in which the ribosome is translating an mRNA sequence. We found the expected tri-nucleotide periodicity consistent with translation in the correct reading frame of the annotated CDSs. These two features were observed in the majority of the 21 families, indicating a robust identification of annotated CDSs in MPRP across different viruses (**Figure S4**).

**Figure 2.**
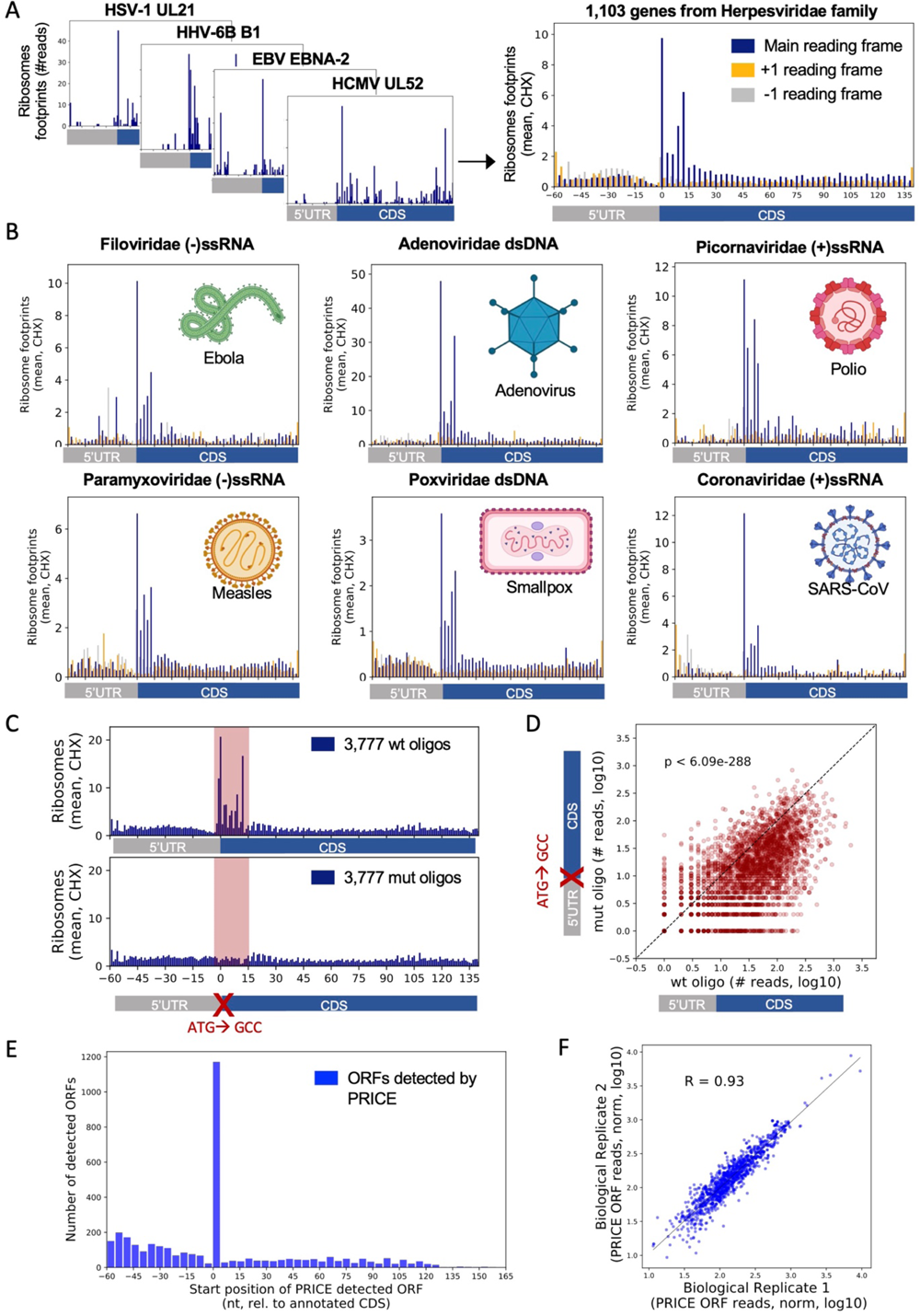
Annotated CDSs measurements and ORF discovery. **(A)** (Left) Example of four individual oligos representing genes from the Herpesviridae family and the number of ribosome footprints observed in the MPRP assay. (Right) Metagene analysis showing the average of ribosome footprints in each position along 1,103 genes from the Herpesviridae family. Different colors represent the three reading frames (blue, 0; orange, +1; gray, -1). **(B)** Average number of ribosome footprints in each position for six additional viral families. A virus representing each family is illustrated in each plot. **(C)** Comparing the average number of ribosome footprints between oligos containing the wt start codon (upper graph) to those in which the annotated start codon was mutated to GCC (lower graph). Shown are the average ribosome footprints in each position across 3,777 oligos. The region with maximum information from footprints containing the wild type or mutated start codon (position -3 to +15) is highlighted in red. **(D)** Pairwise analysis of the number of ribosome footprints on 3,777 viral CDSs with either the wild type start codon or a GCC mutant. Showing the number of footprints in position -3 to +15 relative to the annotated start codon in each oligo. p<10^−288^, Wilcoxon signed-rank test. **(E)** ORF discovery using PRICE. Showing the number of ORFs that were detected in each position (referring to the ORF start position). **(F)** Comparing the number of reads estimated by PRICE for the detected CDSs in two biological replicates. R=0.93, Pearson correlation.

We next confirmed that the detected enrichment of ribosomes in the CDS region resulted from translation initiation at the annotated start codon. We examined ribosome footprints in the presence of an LTM inhibitor, which specifically inhibits initiating 80S complexes with an empty E-site. We found a clear enrichment in the annotated start codon for 19 of the 21 viral families tested (**Figure S5**). We also used reverse genetics to test the contribution of the annotated start codon sequence to CDS translation in the MPRP. We designed oligos with an identical sequence to 3,777 annotated CDSs but mutated the three nucleotides of the start codon to GCC. When comparing ribosome footprints in the region flanking the annotated start codon, we found a substantial reduction in the number of footprints in the mutated oligos (Figure 2C). Since the size of the ribosome footprint is ∼29nt, footprints that do not span the region of three mutated nucleotides are mapped to both the mutated and wt oligo, resulting in similar ribosome occupancy profile downstream of the first few codons (outside the highlighted region in Figure 2C). We then performed pairwise comparison of the number of ribosome footprints for each viral CDS in the wild type and mutated oligo and found a significant reduction in the number of ribosome footprints on the mutated oligos (p<10^-288^, Figure 2D).

We used the annotated CDSs to evaluate the performance of the computational pipeline that we used to infer ORFs in the MPRP experiment. When running PRICE, we did not indicate which viral oligos contain annotated CDSs in order to evaluate their discovery in an unbiased fashion. Rather, we used endogenous ribosome footprints from annotated human CDSs to train PRICE and learn essential parameters of the ribosome footprints in the experiment (see Methods). PRICE has successfully captured 1,136 of 3,976 viral CDSs (28%) initiating at the annotated start codon (false discovery rate 0.05, Figure 2E). The observance of only 28% of the annotated CDSs may represent inherent limitations of the MPRP, assaying the translation of viral sequences in non-infected cells (see Discussions). However, the high specificity of ORF discovery in the expected location along the oligo (Figure 2E) provides high confidence for the thousands of ORFs detected by the assay. We also found high correlation between ribosome occupancy on PRICE-detected CDSs in the two biological replicates (R=0.93, Figure 2F). Together, these data show that MPRP successfully identifies translation initiation and elongation of ribosomes on annotated CDSs across multiple viral families.

### Comparing ORF detection in MPRP and ribosome profiling of HCMV-infected cells

In addition to annotated CDSs, we also estimated the discovery of ORFs by comparing MPRP to traditional ribosome profiling done in the context of native viral infection. As part of the design of the pilot library, we included oligos representing the sequence of 716 ORFs that were identified in HCMV-infected cells by ribosome profiling (Stern-Ginossar et al. 2012), termed here “Ribo-seq ORFs” (Figure 1B).

Similarly to the annotated viral CDS, we found clear enrichment of ribosome footprints along the reported Ribo-seq ORFs with tri-nucleotide periodicity indicating translation in the correct reading frame. Importantly, MPRP detected ribosome footprints on both canonical and non-canonical Ribo-seq ORFs, including ORFs with a non-AUG start codon (CUG, ACG, GUG, AUU, UUG, AUC, AUA, AGG, AAG) (Figures 3A**, 3B**). Moreover, MPRP successfully detected translating ribosomes on short ORFs, in the length of 20 aa or less (Figure 3C).

**Figure 3.**
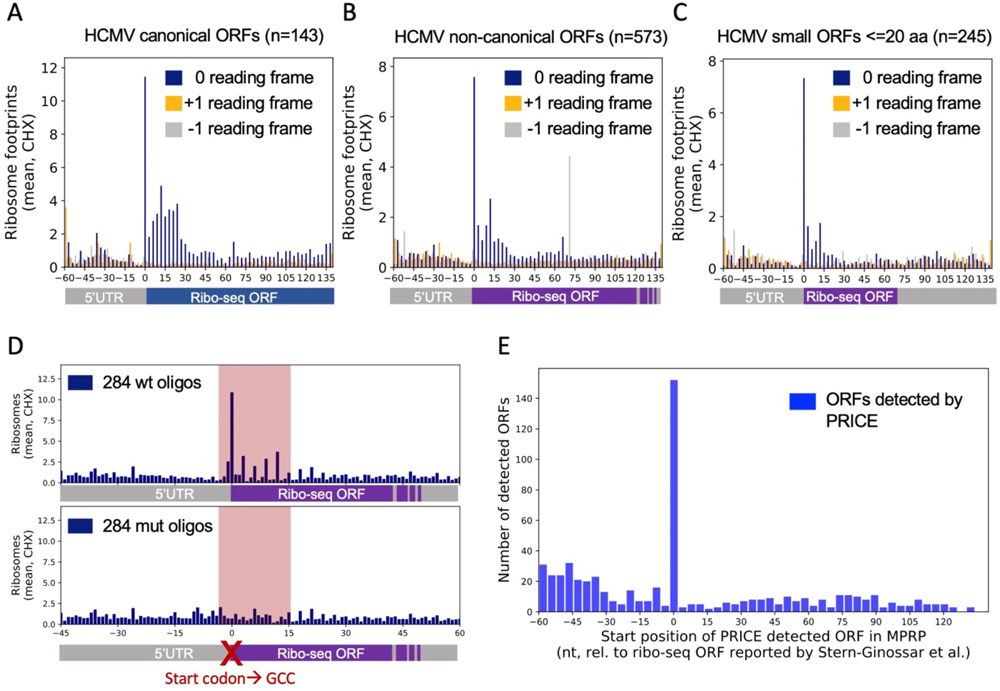
MPRP measurements of canonical and non-canonical ORFs that were identified by traditional ribosome profiling. **(A-C)** Metagene analysis of oligos containing the sequence of ORFs that were identified by ribosome profiling of HCMV-infected cells done by Stern-Ginossar et al. (Stern-Ginossar et al. 2012). Showing the average of ribosome footprints in each position across 143 annotated canonical ORFs (A), 573 non-canonical (B), and 245 non-canonical ORFs in the length of 20 aa or shorter (C). Different colors represent the three reading frames (blue, 0; orange, +1; gray, -1). **(D)** Comparing the average number of ribosome footprints between oligos containing the wt start codon (upper graph) to those in which the reported start codon was mutated to GCC (lower graph). Shown are the average ribosome footprints in each position across 284 oligos, containing Ribo-seq ORFs in the length of 7-45 aa. The region with maximum information from footprints containing the wt or mutated start codon (position -3 to +15) is highlighted in red. **(E)** ORF discovery using PRICE. Showing the number of ORFs that were detected in each position (referring to the ORF start position).

Translation of the Ribo-seq ORFs in the MPRP assay was dependent on the presence of the reported start codon. Mutating the start codon to GCC of 284 Ribo-seq ORFs (in the length of 7-45 aa) resulted in substantial reduction of ribosome footprints compared to wild type oligos (Figure 3D). Unlike annotated CDSs, which mostly initiate with a AUG start codon, the non-canonical Ribo-seq ORFs often initiate from a non-AUG codon. Our findings confirm that the non-AUG codons reported by Stern-Ginossar et al. are essential for translation initiation of the Ribo-seq ORFs and demonstrate how MPRP can be used to functionally characterize the start codon of non-canonical ORFs.

Finally, we ran PRICE to assess the overlap between ORFs that were detected in the original ribosome profiling study and the MPRP experiment. Similarly to the annotated CDS, the observed peak of the PRICE-detected ORF start codon matched the location of the Ribo-seq ORFs in the designed oligo, with 152 of the 716 Ribo-seq ORFs detected (21%) (false discovery rate 0.05, Figure 3E). Altogether, we show that MPRP can capture both canonical and non-canonical ORFs that were identified by traditional ribosome profiling.

### Expanding the repertoire of viral antigens that are presented on the HLA-I complex

Microproteins that are encoded by non-canonical ORFs can contribute to the pool of viral antigens that are presented on the HLA-I complex. Our assay resulted in the discovery of 4,208 non-canonical ORFs, 3,686 of which encode proteins in the length of 8 aa or longer, making them candidates for HLA-I binding.

To assess if these non-canonical ORFs participate in HLA-I presentation, we re-analyzed two HLA-I immunopeptidome datasets from cells that were infected with either HCMV (Erhard et al. 2018) or Vaccinia virus (VACV)(Lorente et al. 2019). In the two studies, the HLA-I complex was immunoprecipitated from infected cells using HLA-specific antibodies and bound peptides were identified using mass spectrometry (MS) (Figure 4A). For the reanalysis, we appended the non-canonical proteins that were detected in HCMV or VACV in the the MPRP assay to the canonical human proteome database and used this combined database to search the raw mass spectrometry files (Erhard et al. 2018; Lorente et al. 2019).

**Figure 4.**
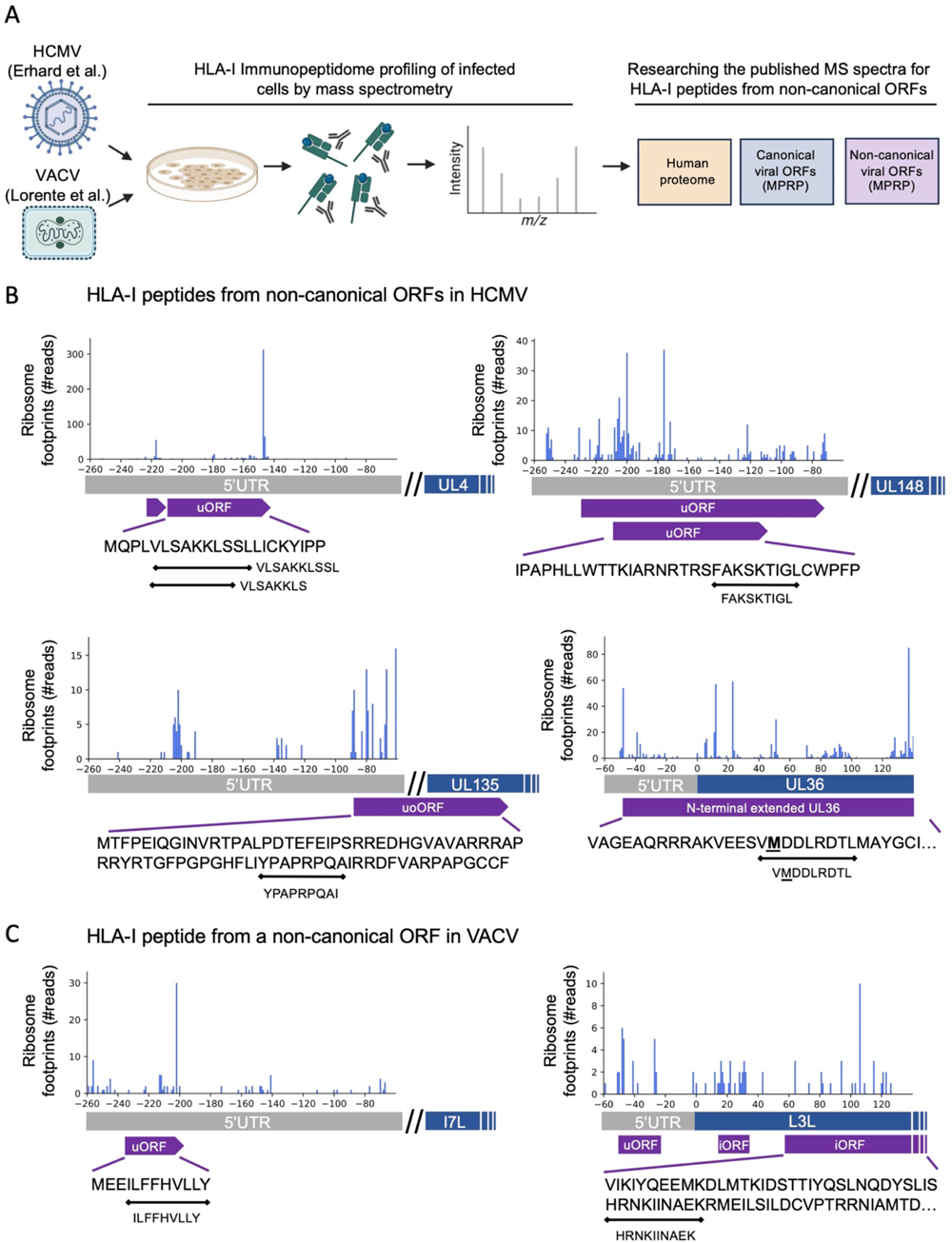
HLA-I peptides derived from non-canonical ORFs in HCMV and VACV. **(A)** (Left) Illustration of the HLA-I immunopeptidome profiling performed by Erhard et al. and Lorente et al. for HCMV-and VACV-infected cells, respectively. (Right) The dataset that we built to research the mass spectrometry raw data including non-canonical ORFs that were identified in the MPRP assay. **(B)** HLA-I peptides that were detected in four non-canonical ORFs of HCMV identified by MPRP: two uORFs in the 5’UTRs of UL4 and UL148, an uoORF in the 5’UTR and coding region of UL135, and an N-extended isoform of UL36. **(C)** HLA-I peptides originating from two non-canonical ORFs in VACV: A uORF in the non-coding region upstream of the I7L coding sequence, and an iORF overlapping the coding region of L3L.

We detected HLA-I peptides that originated from non-canonical ORFs in the HCMV and VACV genomes (Table S2). Overall, we found five potential HLA-I peptides from four non-canonical ORFs in HCMV: an uORF in the 5’UTR of UL4 (VLSAKKLS), and VLSAKKLSSL), an uORF in the 5’UTR of UL148 (FAKSKTIGL), an uoORF in the 5’UTR and coding region of UL135 (YPAPRPQAI), and an N-terminal extended isoform of the UL36 protein (VMDDLRDTL) (Figure 4B). We noted detection of LSAKKLSSL and SAKKLSSL that appear to be in-source fragments of VLSAKKLSSL, and not HLA-I binding peptides, as they elute within 0.1 minutes of each other and do not have strong HLAthena binding predictions. In VACV-infected cells, we found two HLA-I peptides from a uORF in the 5’UTR of I7L (ILFFHVLLY) and from an internal ORF overlapping the coding region of L3L (HRNKIINAEK) (Figure 4C).

Six of the seven detected peptides were predicted as good binders by HLAthena (Sarkizova et al. 2020) (MSi rank <= 2) to at least one of the expressed HLA-I alleles, and all the non-canonical ORFs were supported by a peptide with a good prediction score (Table S2). From HCMV: VLSAKKLSSL(C*0102; MSi rank 0.04), FAKSKTIGL (B*0801; MSi rank 0.02), YPAPRPQAI (B*0801,B*5101; MSi rank 0.5), and VMDDLRDTL (C*0102; MSi rank 2.0). From VACV: ILFFHVLLY (A*0301;MSi rank 0.01) and HRNKIINAEK (B*2705; MSi rank 0.6).

Together, these observations support that non-canonical ORFs identified by MPRP can be translated in the context of native viral infection and that the resulting proteins are presented on the HLA-I complex.

### Exposing uORFs that regulate the translation of canonical viral proteins

In addition to a list of canonical and non-canonical ORFs, our assay provides a detailed view of ribosome footprints across thousands of viral genes that can be harnessed to study translational regulation. Examining the distribution of ribosome footprints in 2,418 viral genes across the 679 human-associated viruses, we identified two main clusters: a group of genes in which most of the footprints were observed in the 5’UTR with low occupancy in the CDS region (5’UTR cluster), and a group of genes in which most of the footprints were detected in the CDS with no evidence for ribosomes in the 5’UTR (CDS cluster) (Figure 5A**, 5B**). The tri-nucleotide periodicity observed in uORFs that were detected in the 5’UTR region indicates that they are actively translated by ribosomes (Figure 5C). Moreover, we found a strong signal of initiating ribosomes at the start codons of these uORFs in the presence of an LTM inhibitor.

**Figure 5.**
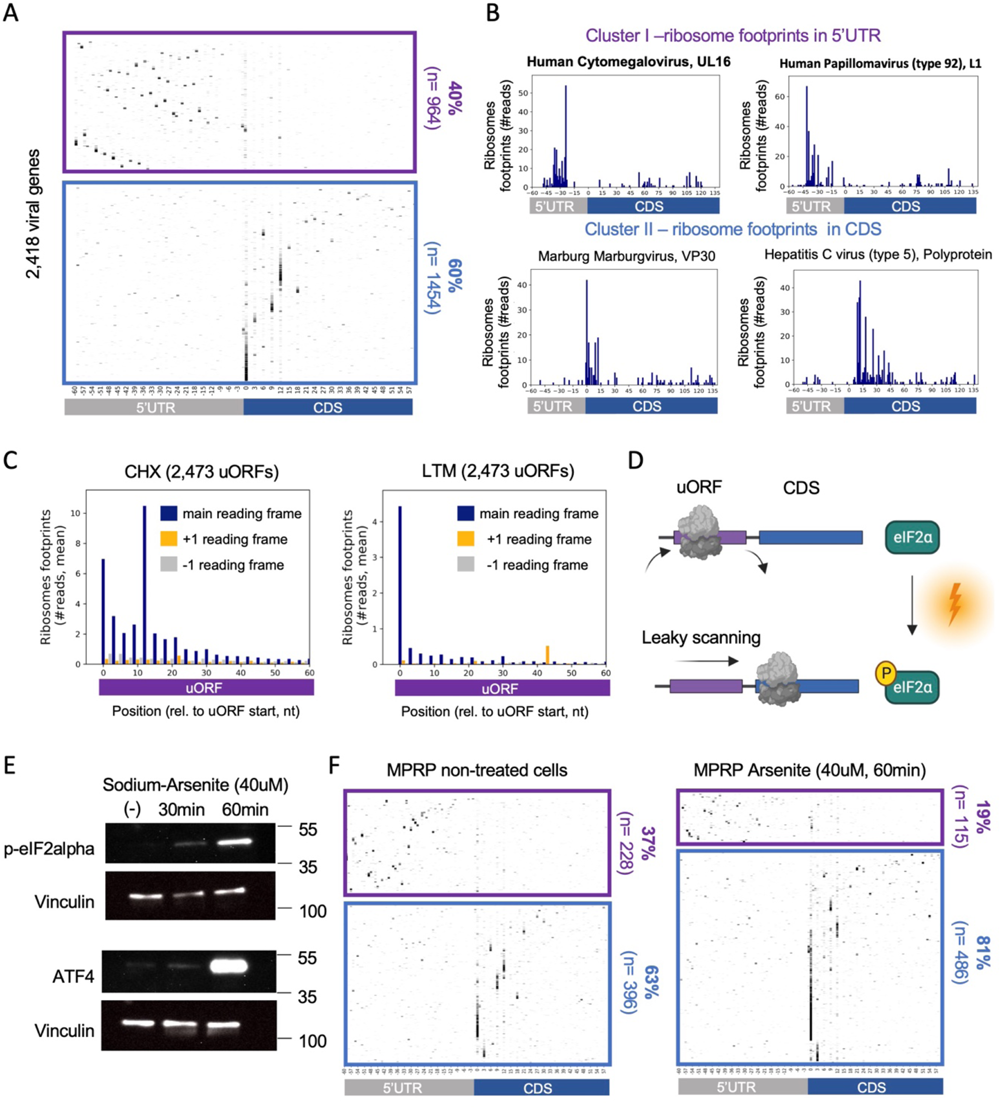
Ribosome densities on uORFs and CDSs in non-stressed cells and in response to eIF2alpha phosphorylation. **(A)** Heatmap showing ribosome footprint densities across 2,418 viral oligos. Each line represents a single viral gene and each column represents the position relative to the annotated start codon. Genes in the upper cluster (purple) had more footprints in the 5’UTR region than the CDS region, and genes in the lower cluster (blue) had more footprints in the CDS than the 5’UTR. **(B)** Example of two individual genes from each cluster and the distribution of ribosome footprints observed in each position. **(C)** Metagene analysis showing the average ribosome footprints in each position along 2,473 uORFs detected by PRICE, relative to the uORF start position. Shown for CHX (Left) and LTM (right) inhibitors. Different colors represent the three reading frames (blue, 0; orange, +1; gray, -1). **(D)** Illustration of uORF-mediated attenuation of translation initiation from canonical CDSs. Ribosomes that initiate at the uORF start codon are less likely to efficiently reinitiate at the downstream start codon of the main CDS. Upon stress, eIF2alpha is phosphorylated as part of integrated stress response, which includes cellular response to viral infection. Ribosomes are more likely to “miss” the uORF start codon and initiate successfully at the start codon of the main CDS. **(E)** Western blot analysis lysates from HEK293T cells treated with 40uM Sodium-Arsenite for 30 and 60 min. Phosphorylated eIF2alpha was detected with a monoclonal phospho S51 antibody (upper panel). ATF4 protein was detected using a polyclonal antibody (lower panel). In both membranes, Vinculin was used as a loading control. **(F)** Repeating the MPRP experiment in HEK293T cells that were treated with 40uM sodium arsenite and in non-treated cells. Shown are heatmaps of ribosome densities across viral oligos and clusters similarly to the analysis in (A).

We hypothesized that viral genes in the 5’UTR cluster have uORFs that attenuate translation initiation from the main CDS (Figure 5D). To test if the translation of the detected uORFs is regulated by cellular stress, we treated cells with sodium arsenite, a potent inducer of eIF2alpha phosphorylation (Andreev et al. 2015). In agreement with Andreev et al., treating HEK293T cells with 40 µM Sodium Arsenite resulted in rapid phosphorylation of eIF2alpha after 60 minutes (Figure 5E). We also observed strong induction of cellular ATF4 protein levels, a well-studied gene that is regulated by uORFs, confirming enhancement of main CDS translation upon arsenite treatment.

We set out to investigate the distribution of ribosome footprints across the pan-viral library in response to arsenite treatment. We transfected HEK293T cells with the library plasmid pool and treated cells with 40 µM sodium-arsenite for 60 minutes prior to ribosome profiling. We found a relative decrease in the fraction of viral genes in which ribosomes were “held” at the 5’UTR (37% to 19% in untreated and treated cells, respectively, Figure 5F) and an increase in the fraction of genes containing ribosome footprints in the CDS (63% to 81% in untreated and treated cells, respectively). This result indicates that in response to eIF2alpha phosphorylation, ribosomes were more likely to bypass the uORFs in the 5’UTR and initiate translation at the main CDS, as expected in the case of inhibitory uORFs. Together, MPRP exposed hundreds of potential uORF that negatively regulate the translation of viral proteins.

## DISCUSSION

We present a method to comprehensively screen the genomes of many viruses for translated regions in a single pooled experiment. Using MPRP, we uncovered thousands of “hidden” ORFs in viral genomes and provided high resolution of ribosome footprints across thousands of annotated viral CDSs.

MPRP has the capacity to identify translated ORFs in a broad spectrum of viruses. Various viral families exhibit unique life cycles, utilizing diverse nucleic acids as their genetic material and employing distinct strategies for replication, RNA transcription, and ribosome recruitment. Furthermore, different viruses replicate within specific subcellular compartments, such as the nucleus, cytoplasm, and viral factories. Despite these variations, we have successfully detected annotated CDSs from many families. Given that all known viruses are unequivocally reliant on the host translation machinery (Stern-Ginossar et al. 2019), their genomes have evolved to be recognized by the 80S ribosome. It is possible that these inherent sequence features facilitate translation initiation when the CDS is expressed from a synthetic construct in uninfected cells.

MPRP’s capabilities hold particular significance in the context of researching highly pathogenic viruses, as our comprehension of their biology is limited due to the scarcity of available high-containment facilities. MPRP, employing 200nt long viral fragments, can be conducted in a BSL-2 setting, rendering it accessible to laboratories worldwide. Our study effectively identified annotated CDSs in families of viruses designated as high-priority pathogens with the potential for outbreaks and global pandemics by the World Health Organization (WHO). Evident signatures of elongating and initiating ribosomes, including tri-nucleotide periodicity, were found in Filoviridae (Ebola virus disease and Marburg virus disease), Paramyxoviridae (Nipah and henipaviral diseases), Arenaviridae (Lassa fever), Coronaviridae (Middle East respiratory syndrome {MERS} and Severe Acute Respiratory Syndrome {SARS}), and Flaviviridae (Zika virus) (Figures 2B**, S4, and S5**). Subsequent MPRP experiments have the potential to deepen our comprehension of the translational regulation and host factor dependencies of these highly pathogenic viruses, which will inform future therapeutics.

Our study unveiled a previously unexplored realm of viral antigens that could contribute to HLA-I presentation and T cell recognition. The identification of peptides in MS experiments relies on a predefined dataset and requires a list of the proteins present in the sample. Consequently, the lack of comprehensive information regarding the complete translatome of viral genomes hinders the detection of HLA-I peptides. It has been suggested that incorporating data from ribosome profiling could enhance the optimization of false-and failed-discovery rates (Holly and Yewdell 2023). Utilizing the list of ORFs detected by our assay, we revealed HLA-I peptides originating from non-canonical ORFs in HCMV and VACV (Figure 4). As researchers generate immunopeptidome data for other viruses, our pan-viral non-canonical ORFs list can serve as a resource for the community, enabling rapid exploration of the new immunopeptidomes. Furthermore, these non-canonical ORFs can also be integrated into the design of peptide pools for T cell assays. While T cell assays almost exclusively assess responses against canonical proteins, we recently demonstrated that some T cell epitopes from non-canonical ORFs induce more potent T cell responses compared to canonical epitopes (Weingarten-Gabbay et al. 2021). Thus, the incorporation of non-canonical ORFs into T cell assays has the potential to enhance their sensitivity and facilitate the identification of vaccine targets.

We reveal an unprecedented number of viral uORFs that likely play a role in gene expression regulation. Proper temporal expression of viral proteins is crucial for their life cycle. For instance, viruses must inhibit host antiviral genes before genome replication to avoid innate immune responses. Structural protein synthesis and virion assembly usually follow sufficient genome replication. Viruses employ ’transcriptional programs’ for ’early,’ ’intermediate,’ and ’late’ genes, often using distinct promoters or controlling transcription levels through the arrangements of genes along their genome. Recent research highlights translation control, and particularly the role of uORFs, in temporal viral gene expression regulation. Enrichment of uORFs in late genes of HHV-6 and HCMV suggests roles in temporal gene control (Finkel et al. 2020). However, unlike transcription-related mechanisms, uORF-mediated gene regulation in viruses remains less understood. Our study uncovers numerous potential uORFs in viral genomes, exhibiting signs of genuine translation, including tri-nucleotide periodicity and enriched initiating ribosomes at uORF start codons (Figure 5C). Notably, many of these uORFs respond to eIF2alpha phosphorylation, resulting in enhanced translation of the primary CDS. Given that eIF2alpha phosphorylation levels rise during infection, the expression of viral genes controlled by these uORFs would be augmented in the later stages of infection. This suggests their function as *cis*-regulatory elements of ’late’ genes.

It is important to acknowledge the limitations inherent in this study. The viral sequences examined here were assessed independently of the broader genome context and were evaluated in non-infected cells. MPRP does not capture the distinctive biology occurring within cells infected by each of the many viruses studied here. Thus, it is plausible that our assay lacks host and/or viral proteins necessary for the translation of certain ORFs. Indeed, MPRP has captured 28% of annotated CDSs and 21% of previously reported non-canonical ORFs. However, it still exposed thousands of new ORFs, providing throughput unparalleled by alternative approaches. While the results from this study should be interpreted with the caution typical of high-throughput screens, there is substantive evidence supporting the identification of genuine ORFs. This evidence arises from the anticipated signature of elongating and initiating ribosomes, the reproducibility of measurements across different cell types, the notable decrease in ribosome footprints upon mutating AUG and non-AUG start codons, the corroborative support from HLA-I peptides identified through mass spectrometry, and the responsiveness of uORFs to stress conditions. In total, our study yields thousands of promising candidates for unexplored ORFs across hundreds of human-associated viruses, which can enhance our understanding of viral biology and contribute to vaccine development.

## METHODS

### Cell culture

Human embryonic kidney HEK293T cells (female) and human lung A549 cells (male) were maintained at 37°C and 5% CO2 in Dulbecco’s modified Eagle’s medium (DMEM) supplemented with 10% fetal bovine serum (FBS) and 1% penicillin and streptomycin.

### Library amplification and cloning

ssDNA oligo libraries were synthesized by Agilent Technologies. Each library was dissolved in 100µL Tris 10mM pH 8 and amplified by PCR with specific primers (Fw: GTGAACCGTCAGATCGCCTCGGCACTCCAGTCCT, Rv: AGAGGGTTAGGGATAGGCTTACCTCAGGCTAGTGCGGACCGAGTCG) using NEBNext Ultra II Q5 HotStart (NEB). PCR cycling parameters were set as follows: initial denaturation at 98°C for 30 seconds, 12 cycles or 98°C for 10 seconds, 63°C for 10 seconds, then 72°C for 10 seconds, followed by a final elongation step at 72°C for 5 minutes. The library oligos were amplified in 8 50µL reactions, pooled together, and concentrated to 70µL using Amicon Ultra-0.5ml centrifugal filters. The PCR product was then purified using AMPure XP beads (Beckman Coulter) at 1.8X concentration. A pcDNA3.4 backbone was amplified with specific primers (Fw: GGTAAGCCTATCCCTAACCCTCT, Rv: AGGCGATCTGACGGTTCAC) using NEBNext Ultra II Q5 HotStart (NEB). The library insert and linearized backbone were assembled using the NEBuilder HiFi DNA assembly master mix and half of the reaction volume was transformed by electroporation into 100µL of NEB 10-beta electrocompetent *E. coli* according to the manufacturer’s instructions. Transformed cells were allowed to recover for 1 hour at 37°C in SOC before being grown for 6.5 hours at 37°C in 100mL LB containing 100µg/mL carbenicillin. A plated dilution series of transformed bacteria was used to verify that the transformation efficiency exceeded 1,000-fold the complexity of the library. Pooled library plasmid was isolated from bacteria using the Qiagen Plasmid Plus Midi Kit.

### Library transfection into cells

#### HEK293T transfection

4 million HEK293T cells were plated in a 10cm tissue culture dish. After 24 hours, cells were transfected with 10µg of library-containing plasmid using X-tremeGENE 9 transfection reagent (Sigma-Aldrich) at a ratio of 3:1 reagent to DNA (30µL transfection reagent for 10µg plasmid) according to the manufacturer’s instructions. Ribosome profiling analysis was performed 24 hours post-transfection.

#### A549 transfection

3 million A549 cells were plated in a 10cm tissue culture dish. After 24 hours, cells were transfected with 15µg of library-containing plasmid using Mirus TransIT-X2 (MIR 6004) transfection reagent at a ratio of 2:1 reagent to DNA (30µL transfection reagent for 15µg plasmid) according to the manufacturer’s instructions. Ribosome profiling analysis was performed 24 hours post-transfection.

### Ribosome Profiling

Ribosome profiling protocol was adapted from McGlincy and Ingolia (McGlincy and Ingolia 2017).

#### Ribosome footprints purification

10cm dishes of library-transfected HEK293T or A549 cells were treated with 100µg/mL cycloheximide (CHX) for 1 min prior lysis. Cells were washed with ice-cold PBS containing 100µg/mL CHX and lysed on ice by scraping in 400ul lysis buffer (20 mM Tris pH 7.4, 150 mM NaCl, 5 mM MgCl_2_, 1 mM DTT, 100µg/mL CHX, 1% Triton X-100, 25 U/mL Turbo DNase). Lysates were incubated on ice for 10 min and triturated cells ten times through a 25 gauge needle. Lysates were centrifuged for 10 min at 20000 x g at 4°C and supernatant was removed into a new tube. RNA concentration was determined using a Qubit RNA BR kit. Lysates were aliquoted to cryotubes and flash frozen in liquid nitrogen. 30µg total RNA was digested with 15 units of RNase I (Epicenter) in 200µl polysome buffer (20 mM Tris pH 7.4, 150 mM NaCl, 5 mM MgCl_2_, 1 mM DTT, 100µg/ml CHX) for 45 min at room temperature. 10uL SUPERase*In were added to stop RNase I digestion and tubes were transferred to ice. Ribosome protected mRNA fragments (RPFs) were enriched using MicroSpin S-400 columns (GE Healthcare, catalog # 27-5140-01). Columns were pre-washed with a 3mL polysome buffer. 100uL of RNAse I digested lysates were loaded to each column and purified RPFs were collected to a new tube by centrifuging the column at 600 x g for 2 min. 400ul TRIzol was added to purified RPFs and RNA was extracted using Direct-zol kit according to manufacturer’s instructions, including on column DNase I treatment, and eluted in 50uL water. RNA was participated by adding 38.5ul water, 1.5uL GlycoBlue, 10uL 3M NaOAc pH 5.5 and 150uL isopropanol and incubating on ice for 1hr (or overnight at −20°C). RNA was pelleted by centrifugation for 30 min at 20,000 x g 4°C, air dried for 10 min and resuspended in 5uL 10mM Tris pH 8. RPFs in the size of 26-34 nt were selected on a 15% polyacrylamide TBE-Urea. RNA was extracted from the gel by adding 400uL RNA gel extraction buffer (300mM NaOAc pH 5.5, 1mM EDTA, and 0.25% SDS), freezing samples for 30 min on dry ice and thawing overnight at room temperature with a gentle mixing nutator. RNA was participated, washed with ice-cold 75% EtOH buffer, air dried and resuspended in 3uL water.

#### rRNA+WPRE depletion

rRNA was depleted using an RNase H-based digestion protocol that our group developed to enhance the metagenomic detection of RNA virus genomes in clinical and biological samples (Matranga et al. 2014; Adiconis et al. 2013). In addition to probes targeting rRNAs, we appended probes tailing the WPRE sequence to remove RPFs originating from the plasmid constant region. rRNA-depleted RNA was eluted in 4uL water.

#### Linker ligation, reverse transcription and cDNA circularization

RNA was dephosphorylated using T4 PNK reaction and ligated to a pre-adenylated linker using T4 RNA ligase. Linkers used for ligation (barcode is highlighted): 5′- /5Phos/NNNNN<u>ATCGT</u>AGATCGGAAGAGCACACGTCTGAA/3ddC/ (NI-810), 5′-/5Phos/NNNNN<u>AGCTA</u>AGATCGGAAGAGCACACGTCTGAA/3ddC//5Phos/NNNNN<u>CGTAA</u>AGATCGGAAGAGCACACGTCTGAA/3ddC/ (NI-811), (NI-812), and 5′- 5′- /5Phos/NNNNN<u>CTAGA</u>AGATCGGAAGAGCACACGTCTGAA/3ddC/ (NI-813). Unligated linkers were depleted by adding 5’-deadenylase and RecJf to the ligation reaction. Ligation products were purified using the Oligo Clean & Concentrator kit and reverse transcribed using protoscript II and RT primer 5′ /5Phos/NNAGATCGGAAGAGCGTCGTGTAGGGAAAGAG/iSp18/GTGACTGGAGTTCAGACGTGTGCTC (NI-802). cDNA was gel purified using 10% polyacrylamide TBE-Urea gel. cDNA was extracted from gel by adding 400uL DNA gel extraction buffer (300mM NaCl, 10mM Tris pH 8, and 1 mM EDTA), freezing samples for 30 min on dry ice and thawing overnight at room temperature with a gentle mixing nutator. cDNA was participated, washed with ice-cold 75% EtOH buffer, air dried and resuspended in 12uL 10mM Tris pH 8. cDNA was circularized using CircLigase II.

#### Library construction PCR and deep sequencing

Sequencing library was PCR amplified from circular DNA using Phusion polymerase and illumina-adaptor containing primers (10-12 cycles). Primers used: Fw: 5′- AATGATACGGCGACCACCGAGATCTACACTCTTTCCCTACACGACGCTC (NI-798), Rv: 5′- CAAGCAGAAGACGGCATACGAGATCGTGATGTGACTGGAGTTCAGACGTGTG (NI-799). Amplified library was purified from E-gel EX Agarose Gels 4% followed by SPRI beads cleaning. Library was quantified using Qubit dsDNA-HS and analyzed on a TapeStation D1000. Pooled libraries were sequenced on NextSeq 550 System using a NextSeq V2.5 High Output 75 cycle kit (illumina, 20024906).

#### Ribosome profiling with LTM inhibitor

For LTM treatment, cells were incubated with 10uM LTM for 30 min prior lysis. LTM was also added to PBS and polysome buffer, and lysis buffer at final concentration of 10uM. The rest of the protocol was identical to ribosome profiling with CHX detailed above.

### Sodium Arsenite treatment and western blot analysis

HEK293T cells were treated with 40uM of sodium arsenate (S7400-100g, Millipore Sigma) for 30 or 60 minutes at 37°C in a dark incubator. For western blot analysis, cells were lysed with a lysis buffer containing 1% SDS, 20mM Tris pH 6.8 and protease inhibitor. Protein quantitation was done using a BCA assay (Pierce BCA protein assay kit, cat #23225) and 10µg of protein was loaded on a gel. After overnight transfer, membranes were incubated with the following antibodies: Rabbit anti-ATF4 (Proteintech 10835-1-AP) at dilution of 1:1,000, Rabbit anti-phospho eIF2alpha (abcam ab32157) at dilution of 1:1,000, and Rabbit anti-Vinculin (CST 13901s) at dilution of 1:1,000.

### Mapping ribosomes footprints to the synthetic library

Illumina sequencing reads were pre-processed prior alignment using FASTX-Toolkit. Illumina adapters were removed using fastx_clipper, reads were split according to samples barcodes using fastx_barcode_splitter.pl and UMI sequence was trimmed using fastx_trimmer. Reads that were mapped to rRNA were removed using bowtie (version 1.2.2). Remaining reads were aligned to an artificial genome composed of all library oligos using bowtie.

### ORF discovery using PRICE

We used PRICE to identify ORFs from deep sequencing reads (Erhard et al. 2018). PRICE algorithm requires a predefined set of annotated CDSs in order to estimate the codons that have generated the observed ribosome footprints. When providing the reference genome of the synthetic library, we did not indicate which oligo contains an annotated viral CDS, because we used this information to estimate ORF discovery rate. Instead, we used annotated human CDSs that were translated in library-transfected cells and thus, were exposed to the exact same experimental conditions (e.g. RNase I concentration and incubation time that can impact footprints size). We generated a chimeric reference genome composed of chromosome 1 in hg19 and an artificial chromosome composed of 15,000 oligos of the pan-viral library. In addition, we generated a gtf file with the annotations of hg19 and library oligos required for PRICE predictions. For each experiment, we mapped deep sequencing reads to the chimeric fasta file using Bowtie. We ran PRICE with bam files from all the experiments done in HEK293T cells. We filtered ORFs a p-value ≤0.05 after correcting for false discovery rate (FDR). ORFs were then defined by extending each initiating codon to the next in-frame stop codon in the corresponding viral genome.

### Tri-nucleotide periodicity analysis

To determine the reading frame, we used ribosome footprints in the exact length of 29nt and plotted the position of the first nt of the sequencing read. Positions were corrected for the P site offset (12nt).

### Re-analysis of HLA-I immunopeptidome datasets

MS/MS spectra from publically available immunopeptidomics data were interpreted using Spectrum Mill (SM) v 07.11.216 (proteomics.broadinstitute.org).

Using the SM Data Extractor module for HLA-I immunopeptidomes, spectral merging was disabled, the precursor MH + inclusion range was 600–3000, and the spectral quality filter was a sequence tag length >1 (i.e., minimum of three peaks separated by the in-chain masses of two consecutive amino acids).

Parameters for the SMMS/MS search module for HLA-I immunopeptidomes reported in(Lorente et al. 2019), included: no enzyme specificity; precursor and product mass tolerance of ±10 ppm; minimum matched peak intensity of 30%; ESI-ORBITRAP-CID-HLA-v3 scoring; fixed modification: cysteinylation of cysteine; variable modifications: oxidation of methionine, deamidation of asparagine, acetylation of protein N-termini, and pyroglutamic acid at peptide N-terminal glutamine; and precursor mass shift range of −18 to 136 Da. MS/MS spectra were searched against a protein sequence database that contained 310566 entries, including all University of California Santa Cruz Genome Browser genes with hg19 annotation of the genome and its protein-coding transcripts (63,691 entries), 602 common laboratory contaminants, 2043 curated small ORFs (lncRNA and upstream ORFs [uORFs]), 237,427 novel unannotated ORFs (nuORFs) supported by ribosomal profiling nuORF DB v1.037, as well as translated canonical and noncanonical ORFs >7 amino acids observed in the 15K ribosomal profiling library experiment mapping to Cytomegalovirus.

Parameters for the SM MS/MS search module for HLA-I immunopeptidomes reported in Erhard et al., (Erhard et al. 2018) included: no enzyme specificity; precursor and product mass tolerance of ±10 ppm; minimum matched peak intensity of 30%; ESI-QEXACTIVE-HCD-HLA-v3 scoring; variable modifications: cysteinylation of cysteine; oxidation of methionine, deamidation of asparagine, acetylation of protein N-termini, and pyroglutamic acid at peptide N-terminal glutamine; and precursor mass shift range of −18 to 136 Da. MS/MS spectra were searched against a protein sequence database that contained 311427 entries, including all University of California Santa Cruz Genome Browser genes with hg19 annotation of the genome and its protein-coding transcripts (63,691 entries), 602 common laboratory contaminants, 2043 curated small ORFs (lncRNA and upstream ORFs [uORFs]), 237,427 novel unannotated human ORFs (nuORFs) supported by ribosomal profiling nuORF DB v1.037(Ouspenskaia et al. 2022), as well as translated canonical and noncanonical ORFs >7 amino acids observed in the 15K ribosomal profiling library experiment mapping to Alphavirus Orthopoxvirus.

Using the SM Autovalidation module, peptide-spectrum matches (PSMs) for individual spectra were confidently assigned by applying target-decoy based FDR estimation to achieve <1.0% FDR. For HLA-I immunopeptidomes, PSM-level thresholding was done with a minimum peptide length of 7, minimum backbone cleavage score of 5, and <1.0% FDR across multiple replicates is present. Allowed precursor charges were HLA-I: 1–4. PSMs mapping to non-canonical ORFs were filtered for 8-11mers to perform HLAthena predictions and manually inspected to exclude peptides that were a result of in-source fragmentation (peptides within nested sets with retention times <20 secs apart).

### Subset-specific FDR filtering for nuORFs

Subset-specific FDR was performed as previously described (Ouspenskaia et al. 2022). The aggregate FDR was set to <1% as described above. FDR for the subset of nuORF peptides requires more stringent score thresholding to reach a suitable subset specific FDR of <1%. Subsets of nuORF types were thresholded independently from the HLA dataset through a two step approach. First, PSM scoring metrics thresholds were tightened on the nuORF subset: minimum SM score of 7, minimum percent scored peak intensity (SPI) of 50%, precursor mass error of ±5 ppm, minimum backbone cleavage score (BCS) of 5. This allows nuORF distributions for each metric to meet or exceed the aggregate distributions. Second, remaining nuORF type subsets with FDR estimate above 1% were further subjected to a grid search to determine the lowest values of BCS and SM score that improved the FDR to <1% for each ORF type in the dataset.

### HLAthena HLA-I peptide presentation predictions

HLA peptide prediction was performed using HLAthena (Sarkizova et al. 2020). For HCMV infected HF99-7, HLA A*01:01, A*03:01, B*08:01, -B*51:01, C*07:01, C*01:02 were used for HLAthena predictions (Erhard et al. 2018). For VACV-infected HOM-2 cells, HLA A*03:01, B*27:05:02, and C*01:02 were used for HLAthena predictions(Lorente et al. 2019).

## ACKNOWLEDGMENTS

We thank Tamara Ouspenskaia, Steven Reilly, James Xue, Aaron Lin, Catherine Freije and Yingpu Yu for many valuable discussions. We thank Nicholas McGlincy and Nicholas Ingolia for sharing their detailed ribosome profiling protocol with the broad community of researchers (McGlincy and Ingolia 2017), which contributed to the development of the MPRP. This study was supported in part by grants from the National Institute of Allergy and Infectious Diseases (U19AI110818 to P.C.S.) and the United States Department of Agriculture (58-3022-2-031 to P.C.S). This work was supported in part by grants P01CA206978 to S.A.C from the NIH, and grants U24CA270823, U01CA271402 to S.A.C. from National Cancer Institute (NCI) Clinical Proteomic Tumor Analysis Consortium program. S.W.-G. is the recipient of a Human Frontier Science Program fellowship (LT-000396/2018), EMBO non-stipendiary long-term fellowship (ALTF 883-2017), the Gruss-Lipper postdoctoral fellowship, the Zuckerman STEM Leadership Program fellowship, and the Rothschild Postdoctoral Fellowship.

## TABLE LEGENDS

**Table S1. ORFs detected in MPRP measurements of the pan-viral library**

**Table S2. HLA-I peptides originating from non-canonical ORFs in HCMV and VACV**

