## Supplemental Figures for "Pan-viral ORFs discovery using Massively Parallel Ribosome Profiling"

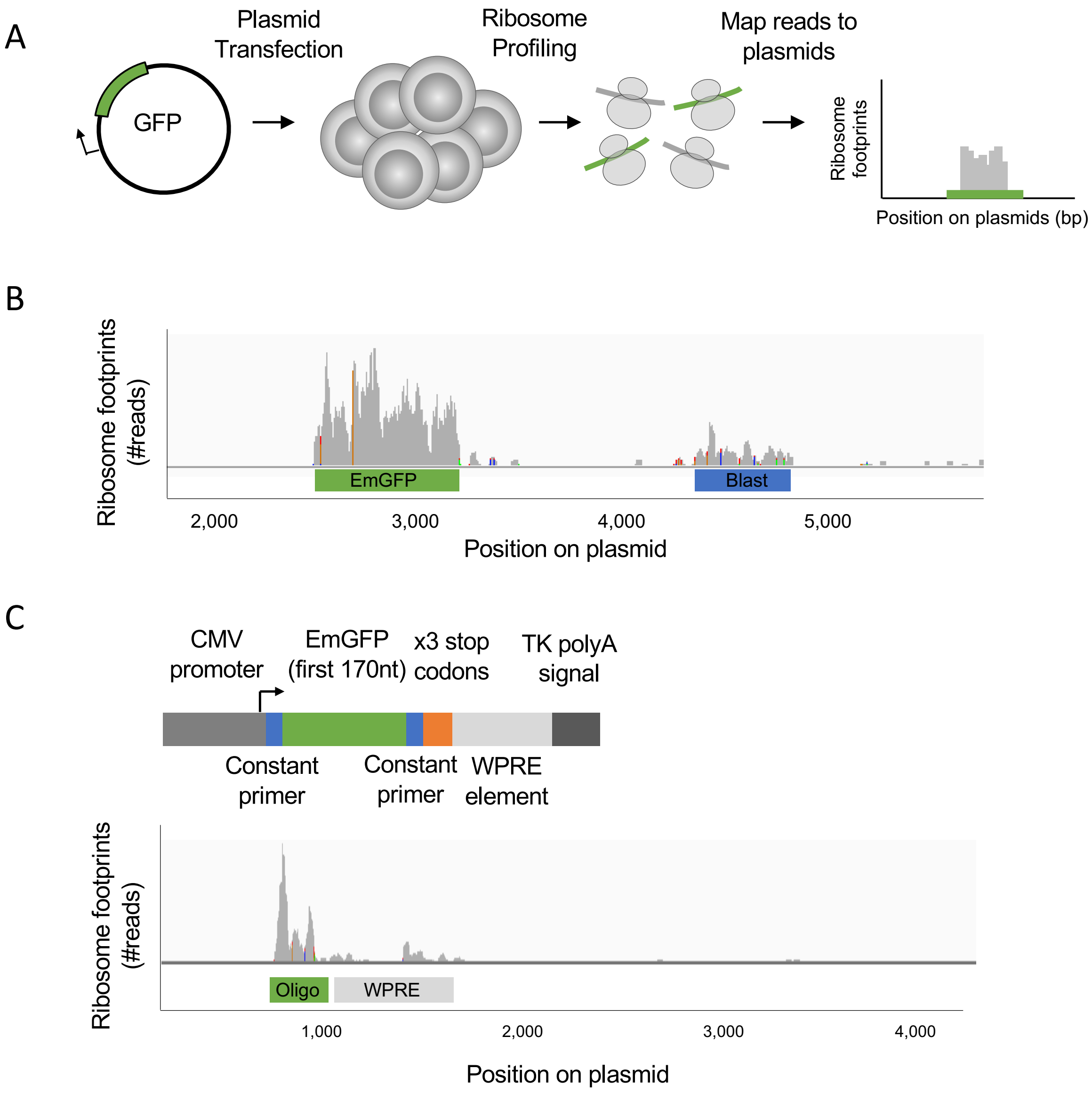


**Figure S1. Ribosome profiling of exogenous ORF from overexpression plasmid**.
**(A)** Illustration of EmGFP plasmid transfection followed by ribosome profiling. **(B)** Mapping ribosome footprints to EmGFP plasmids using Integrative Genomics Viewer (IGV). **(C)** (Top) The design of a truncated EmGFP construct mimicking the synthetic library oligos including the constant primers and the plasmid cloning site. (Bottom) Mapping ribosome footprints to the truncated EmGFP oligo using IGV.

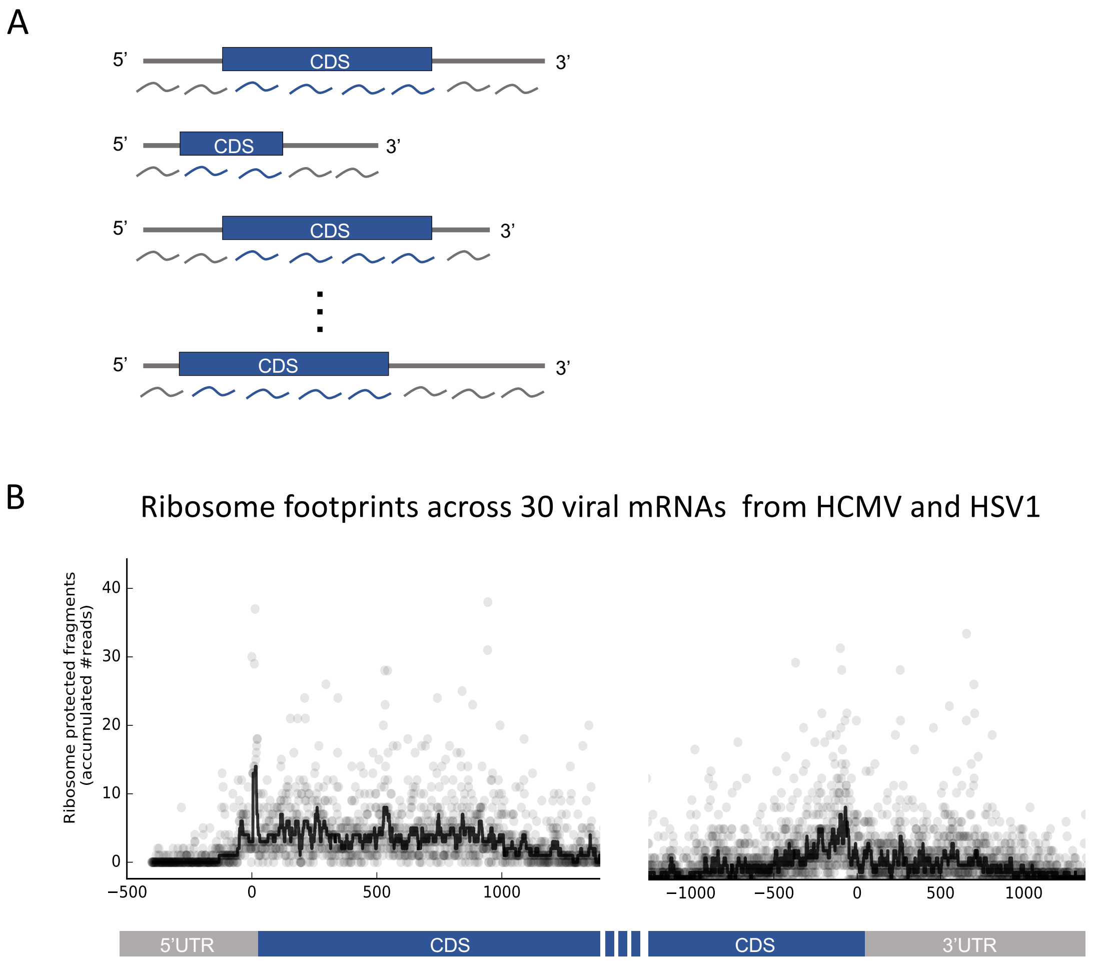


**Figure S2. MPRP measurements of tailing oligos across 30 transcripts of HCMV and HSV-1**
**(A)** The design of tailing oligo across 30 mRNAs. **(B)** Showing the total number of ribosome footprints in each position relative to the CDS start codon or the stop codon in 30 annotated mRNAs (14 of HSV-1 and 16 of HCMV).

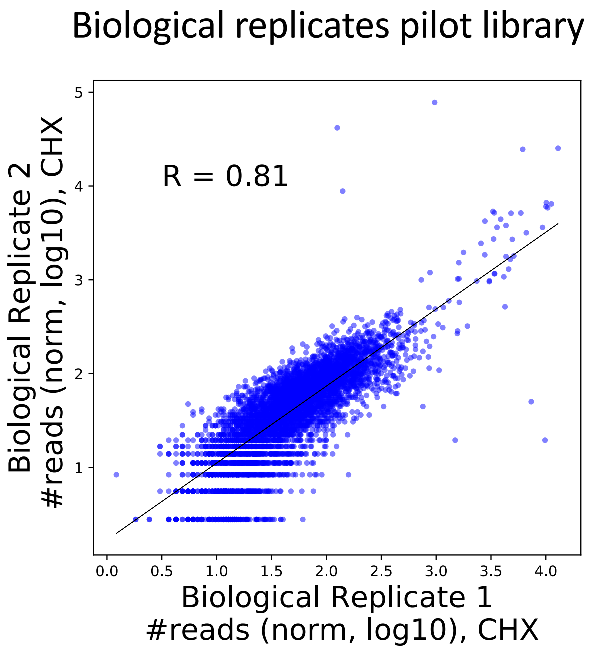


**Figure S3. Biological replicate of MPRP measurements of the pilot library**

Comparing the number of ribosome footprints mapped to 5,170 oligos of the pilot library in two biological replicates of MPRP experiment in HEK293T cells. R=0.81, Pearson correlation.

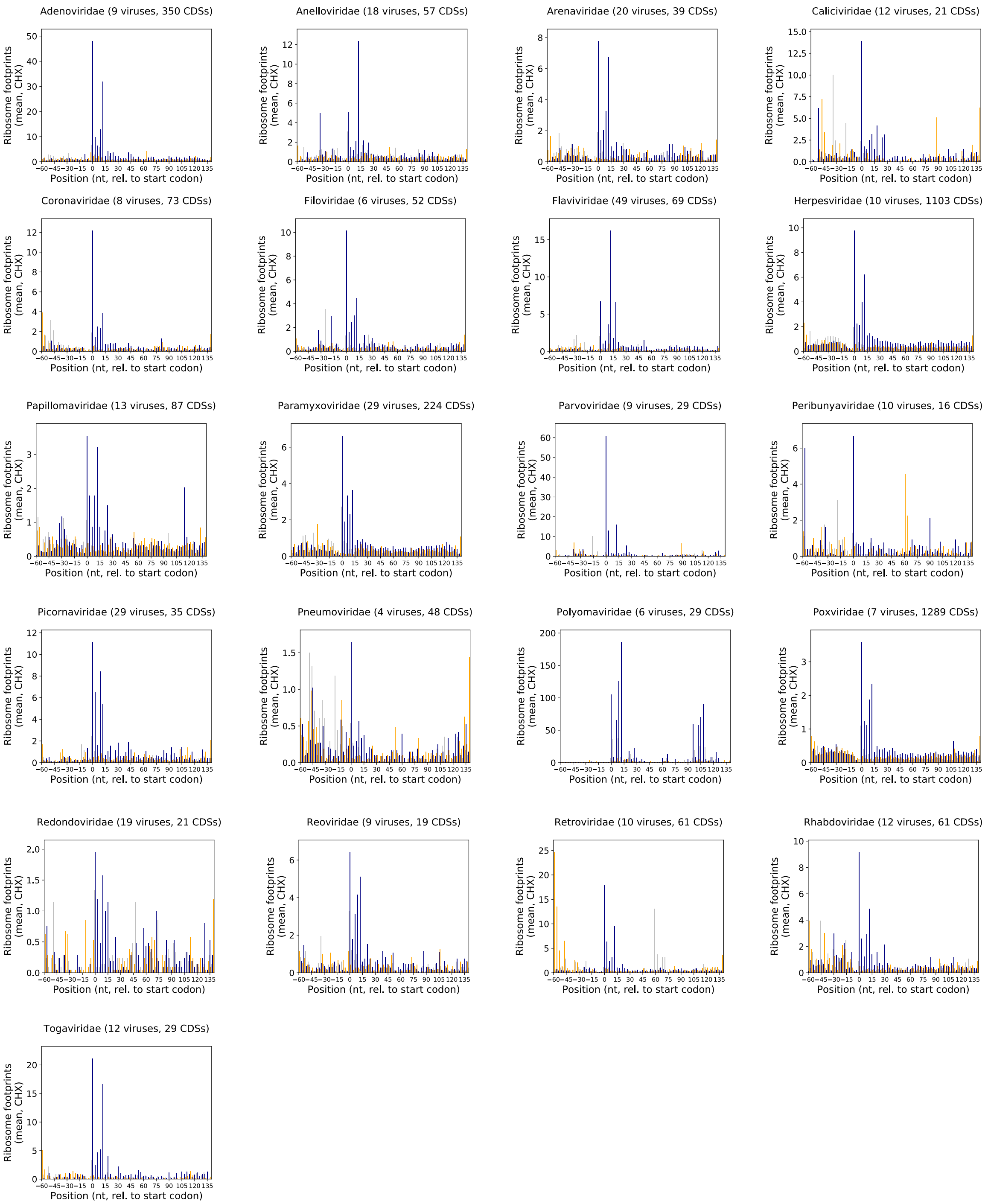


**Figure S4. Metagene analysis of elongating ribosome footprints across 21 viral families**

Average number of ribosome footprints in each position for 21 viral families observed in MPRP measurements after treatment with CHX to inhibit elongation ribosome. Different colors represent the three reading frames (blue, 0; orange, +1; gray, -1)

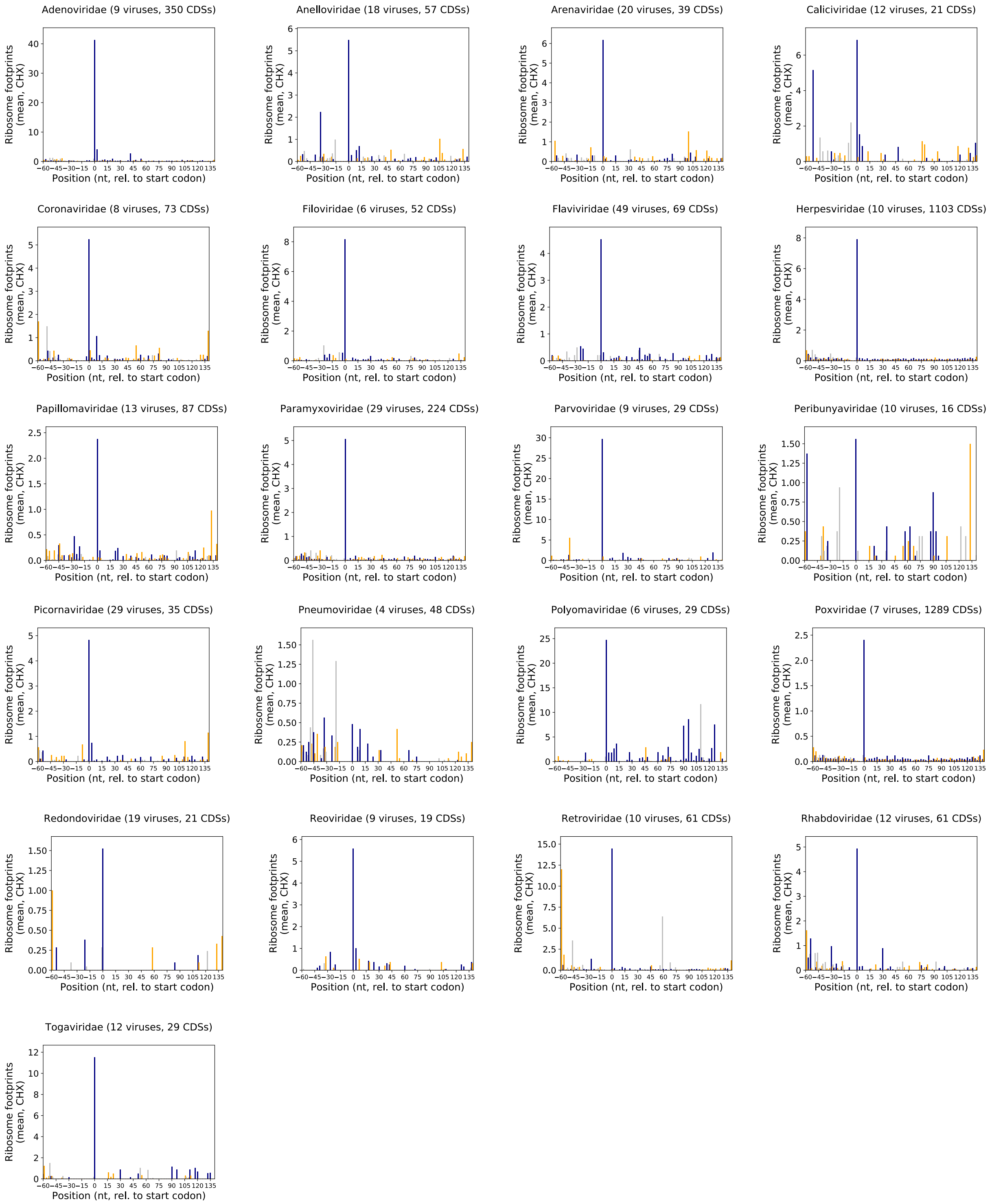


**Figure S5. Metagene analysis of initiating ribosome footprints across 21 viral families**
Average number of ribosome footprints in each position for 21 viral families observed in MPRP measurements after treatment with LTM to inhibit initiating ribosome. Different colors represent the three reading frames (blue, 0; orange, +1; gray, -1)
